# A global synthesis of yeast in microbiomes

**DOI:** 10.64898/2025.12.17.695015

**Authors:** Chieh-Ping Lin, Alberto Geroldi, Nelly Selem, Gianni Liti, Isheng Jason Tsai

## Abstract

Yeasts are widespread members of microbial communities across terrestrial, aquatic, and host-associated environments, yet they remain underrepresented in microbiome studies due to low abundance and methodological biases. By combining a literature review with a meta-analysis of ∼44,000 fungal metabarcoding samples from the GlobalFungi database, we show that yeasts occur in over 90% of samples, confirming their global ubiquity. Basidiomycetous lineages—especially *Agaricomycotina*—were most frequently detected, whereas *Saccharomycotina* were more restricted to anthropogenic and aquatic settings. Although yeasts typically comprised only ∼0.1% of fungal reads, their distributions were non-random and reflected distinct habitat preferences across environments. In ∼3% of samples, yeasts exceeded 25% of reads, with genera such as *Aureobasidium*, *Hanseniaspora*, and *Saccharomyces* episodically dominating nutrient-rich or human-influenced environments. Cosmopolitan genera including *Vishniacozyma, Solicoccozyma,* and *Rhodotorula* were broadly distributed but remain underreported in microbiome surveys. Shotgun metagenomic data further confirmed yeast presence across diverse microbiomes, though with consistently low coverage, reflecting the “curse of low abundance.” Despite their rarity, yeasts likely play disproportionate roles in nutrient cycling, plant growth, and host interactions. We recommend inclusive multi-kingdom approaches—improved primers, long-read sequencing, and quantitative tools—to better integrate yeasts in microbiomes.

## Introduction

Microbial communities play crucial roles in shaping ecosystem functions, from carbon cycling and nutrient transformation to host health and disease resistance. Over the past two decades, microbiome research has expanded rapidly, but its primary focus has been on bacterial and archaeal components, often overlooking the contributions of eukaryotic microorganisms. This bias is reflected in the dominant use of 16S rRNA gene sequencing, which cannot capture fungal diversity, and in the limited fungal-targeted sequencing within major microbiome projects (Leho Tedersoo et al., 2014; Thompson et al., 2017). Yet, fungi—and yeasts in particular—are indispensable members of microbial consortia in terrestrial, aquatic, and host-associated environments (Boekhout et al., 2021).

Yeasts are unicellular fungi within the Ascomycota and Basidiomycota phyla, exhibiting remarkable ecological breadth—from soils and plant surfaces to aquatic systems and animal microbiomes (Yurkov, 2018). Their metabolic flexibility underpins life strategies ranging from saprophytic and commensal to pathogenic and mutualistic (Buzzini, Lachance, & Yurkov, 2017). In soils, yeasts contribute to nutrient cycling and decomposition; on leaf surfaces, they form part of the phyllosphere community; and in marine systems, they often associate with particles or algal hosts, playing roles in organic matter remineralisation (Bass et al., 2007; Heitman & Amend, 2014). Certain taxa are also notorious pathogens: *Candida albicans* is a major cause of candidiasis in humans, *Cryptococcus neoformans* causes life-threatening meningitis, and *Malassezia furfur* can induce dermatitis (Boekhout et al., 2021; Underhill & Iliev, 2014). Others establish mutualistic associations—for example, basidiomycetous Cystobasidiomycetes that persist as consistent lichen cortical symbionts, a partnership long overlooked in classical fungal ecology (Spribille et al., 2016). Across host-associated niches including the gut, skin, and oral microbiomes, genera such as *Candida* and *Malassezia* interact with host immunity and competing microbes in ways that can be either beneficial or pathogenic depending on context (Underhill & Iliev, 2014).

Despite their functional diversity and ubiquity, yeasts remain underrepresented in microbiome analyses. Standard 16S amplicon surveys target bacteria and archaea, leaving fungi invisible unless internal transcribed spacer (ITS) sequencing or related approaches are applied. Even then, yeasts are often missed due to low relative abundance (the “rare biosphere”), DNA extraction challenges, primer bias, and stringent bioinformatic filters (Aziz et al., 2015; Lynch & Neufeld, 2015; Renzi, Nenciarini, Bacci, & Cavalieri, 2024; Leho Tedersoo et al., 2015). Fungal reference databases also remain less comprehensive than bacterial ones, limiting accurate taxonomic resolution (Kõljalg et al., 2013). These methodological issues have created persistent gaps in our understanding of yeast distribution and ecological function.

While often overlooked in broad surveys, yeasts are well known to dominate in certain specialised contexts. Human-associated niches provide classic examples: *Malassezia* species thrive as skin commensals (Theelen et al., 2018) and *Candida* species regularly dominating mucosal microbiomes (Lai et al., 2023; Nash et al., 2017). In anthropogenic environments, the fermentative yeast *Saccharomyces cerevisiae* is notably dominant in traditional food and beverage fermentations such as bread, beer, wine, and sourdough (Carlino et al., 2024; Dashko, Zhou, Compagno, & Piškur, 2014; Landis et al., 2021). These familiar cases highlight the potential of yeasts to dominate nutrient-rich or host-associated conditions, but they represent only a narrow subset of yeast ecological strategies (Opulente et al., 2024).

Beyond these classical niches, yeasts contribute to a broad spectrum of ecosystems, often at lower abundances. In extreme habitats such as semi-arid soils and deserts, they exhibit resilience to high salinity and temperature (Araya et al., 2023; Segal-Kischinevzky et al., 2022). Aquatic systems, including freshwater and marine habitats, harbor significant fungal diversity, with yeasts actively participating in organic matter remineralization and carbon cycling (Gladfelter, James, & Amend, 2019). Agricultural ecosystems similarly sustain diverse yeast communities shaped by soil chemistry, plant age, and complex microbial interactions (S. Zhu et al., 2023). Even in the gut and oral cavity, where they are rare compared to bacteria, yeasts can influence host health—for example, in metabolic dysfunction-associated fatty liver disease (Zeng & Schnabl, 2024). In forests, especially broadleaf Fagaceae ecosystems, yeasts coexist with diverse microbial consortia and partition niches across leaves, twigs, litter, and soils (Lin et al., 2025; Masinova et al., 2017). Our own work in Taiwan’s subtropical forests, which combine high plant endemism with fungal richness, has shown substantial variation in yeast occurrence and turnover, complementing global perspectives on yeast biogeography (Lin et al., 2025; Mozzachiodi et al., 2022).

In this study, we present a comprehensive synthesis of yeast occurrences across global microbiomes, integrating publicly available datasets including GlobalFungi (Větrovský et al., 2020) and literature searches. We analyse how yeasts vary across habitats, their taxonomic composition, and the challenges that limit their detection. By highlighting the ecological significance of yeasts—even as a numerical minority—we aim to advance the integration of fungal components in microbiome science and emphasise the need for more inclusive, multi-kingdom approaches.

## Results

### Representation of yeast genera across published microbiome studies

We systematically assessed yeast representation across 39 published microbiome studies spanning diverse habitats—including terrestrial, aquatic, and host-associated environments—and employing a range of analytical approaches (isolation, amplicon sequencing, and shotgun metagenomics) with different genetic markers (ITS1, ITS2, LSU) (**Fig. 1a, Supplementary Fig. 1 and Supplementary Table 1**). This survey revealed marked variability in reporting both frequency and dominance of yeasts across studies. Such heterogeneity likely reflects not only genuine ecological differences among environments but also methodological inconsistencies in how abundant or prevalent yeast is defined and reported.

**Figure 1.**
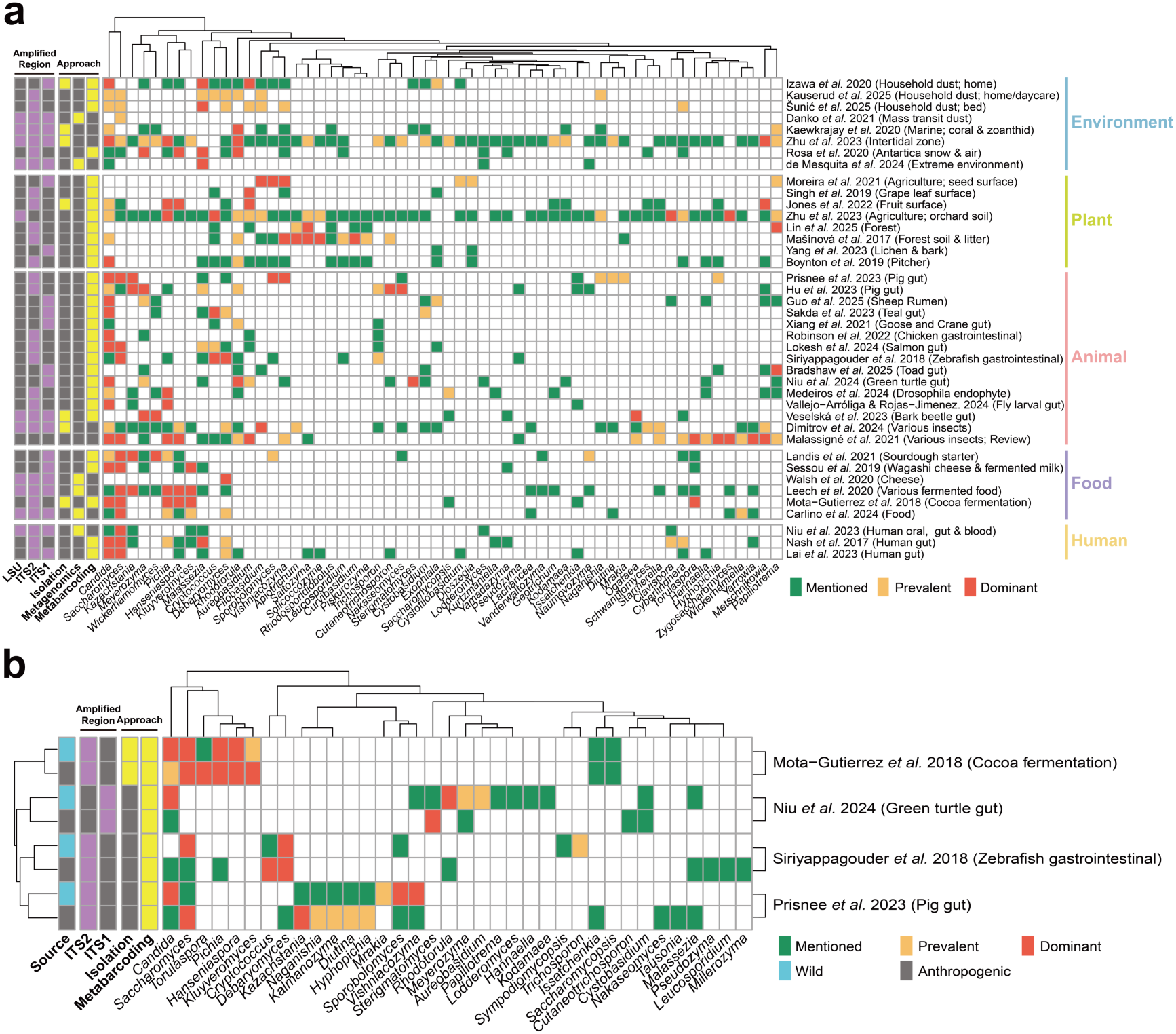
Representation of yeast genera across published microbiome studies. **(a)** Heatmap showing yeast genera that were mentioned (green), detected (yellow), or reported as dominant (red) across 40 representative microbiome studies spanning terrestrial, aquatic, host-associated, and anthropogenic environments. Only genera present in at least four studies are shown. Purple and yellow bars indicate the amplified region and analytical approach used in each study, respectively. Detailed study information is provided in **Supplementary Table S1**. **(b)** Heatmap of four selected studies highlighting the enrichment of particular yeast genera in contrasting wild (blue) versus anthropogenic (grey) environment. More detailed information is shown in **Supplementary Fig. 1** and **Table 1**.

Several genera such as *Candida, Saccharomyces, Hanseniaspora, Pichia, Debaryomyces,* and *Cyberlindnera* were repeatedly emphasised across studies, reflecting their established significance in clinical, fermentative, or host-associated contexts. Likewise, cosmopolitan taxa such as *Meyerozyma* and *Rhodotorula* were frequently observed, underscoring their generalist strategies and broad adaptability. Taxa traditionally known to associate with animal hosts are also frequently detected in a wide range of non-host environments: for instance, *Malassezia* was detected not only in human and animal niches but also in Antarctic snow and air (L. H. Rosa et al., 2020), orchard soils (S. Zhu et al., 2023) and marine ecosystems (Steinbach et al., 2023). Similarly, *Saccharomyces*, long viewed through the lens of domestication in baking and brewing, was identified in soil, bark, wild fruits, and insect samples (Lee et al., 2022; Mozzachiodi et al., 2022), emphasising its broader ecological plasticity. In contrast, genera such as *Tausonia, Glaciozyma,* and *Curvibasidium* appeared only sporadically, suggesting their ecological roles are underrepresented in current microbiome literature. In addition, overlapping sets of genera were reported in intertidal sediments (H.-Y. Zhu et al., 2023) and orchard soil samples (S. Zhu et al., 2023), hinting at shared selective pressures—such as fluctuating moisture, nutrient enrichment, or recurrent microbial interactions—that favour certain adaptable yeasts.

To extend these genus-level comparisons, we assessed whether particular yeast taxa show consistent enrichment patterns across different environmental contexts (**Fig. 1b**). This analysis revealed emerging patterns of divergence between wild habitats and anthropogenic environments shaped by farming, food production, or laboratory conditions. For instance, *Saccharomyces cerevisiae* and *Hanseniaspora uvarum* were repeatedly enriched in fermentation-associated or agricultural niches (Leech et al., 2020; Mota-Gutierrez et al., 2018; Sessou et al., 2019), while *Candida albicans* and *Malassezia restricta* predominated in host-associated microbiomes (Hu et al., 2023; Izawa et al., 2020; Lokesh, Siriyappagouder, & Fernandes, 2024; Nash et al., 2017). By contrast, wild habitats tended to feature a broader array of basidiomycetous and ascomycetous yeasts that were rarely dominant but consistently present. Similar contrasts have been documented in recent targeted studies: in pigs, *Kazachstania slooffiae* was more abundant in intensively raised herds than in feral populations, which harboured a more diverse mycobiome with genera such as *Vishniacozyma* and *Candida* (Prisnee et al., 2023); and in zebrafish, wild individuals hosted mycobiota dominated by Dothideomycetes, while laboratory-reared counterparts were enriched for Saccharomycetes and opportunistic yeasts such as *Debaryomyces* (Siriyappagouder et al., 2018). These cross-system patterns suggest that anthropogenic disturbance funnels yeast communities toward a narrow set of ecologically plastic taxa, whereas wild environments sustain a broader but less frequently studied array of lineages.

### Global yeast occurrences across diverse environments

Our literature survey underlines a bias toward a few well-known yeasts and hints at overlooked diversity. Hence, we surveyed global yeast occurrences using an updated release of the GlobalFungi (Větrovský et al., 2020), which includes 84,972 ITS barcode sequence derived from a broad array of biomes world-wide (**Fig. 2**). Sequencing depth had a pronounced effect on yeast detection. Samples with higher sequencing depths were significantly more likely to detect yeasts (median number of reads: 17,641 sequences with yeasts versus 294 without; **Fig. 2a**, Wilcoxon rank sum test, P<0.0001). Therefore, we restricted downstream analyses to samples with >10,000 reads to ensure adequate detection sensitivity. Across broad environmental categories, yeasts were detected in the vast majority of samples, with high occurrence rates ranging from 87.9% to 98.8% (**Fig. 2b**). Interestingly, substantial differences emerged when considering specific sample types within these broader environments (**Fig. 2c**). Samples associated with lichen and moss samples exhibited much lower yeast detection frequencies. The dominant lichen symbiont fungi (averaging ∼83% of reads) likely outcompete yeasts (Yang, Woo, Oh, Kim, & Hur, 2023). In addition, standard ITS primers may under-amplify basidiomycetous yeasts commonly found in lichens, such as Cystobasidiomycetes and Tremellomycetes, further reducing their apparent detection (Bellemain et al., 2010; Neilan et al., 2014). At a global scale, spatial analysis revealed that yeast sequences were distributed across all continents and major biogeographic zones, underscoring their widespread ecological versatility (**Fig. 2d**).

**Figure 2.**
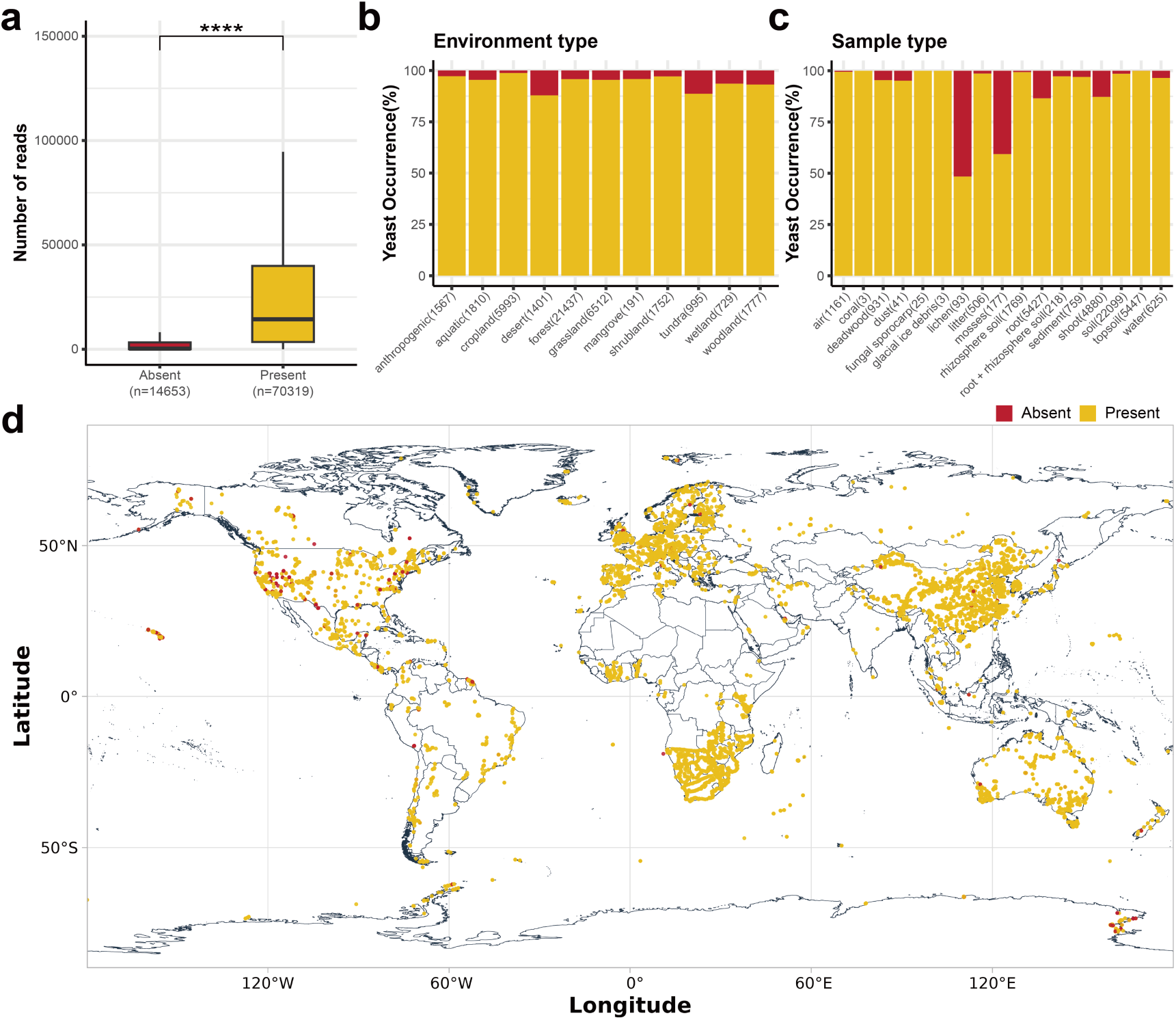
Overview of yeast occurrence across global microbiome samples. **(a)** Boxplot comparing sequencing depth between samples with and without detected yeast taxa. Outliers are excluded. Number in brackets denote number of samples. **(b)** Barplot showing the proportion of samples with yeast presence or absence across major environment types. **(c)** Barplot showing yeast presence or absence across different sample types. **(d)** World map indicating geographic locations of samples, with points colored by yeast detection status. Presence and absence are indicated in yellow and red, respectively.

Subphylum-level analyses further delineated the structure and breadth of yeast communities across environments. Basidiomycetous lineages typically dominated, with Agaricomycotina (average detection rate 89.2%) and Pucciniomycotina (70.9%) representing the most consistently detected subphyla across sample types, followed by Saccharomycotina (60.5%), Pezizomycotina (36.1%) and Ustilaginomycotina (30.0%). Taphrinomycotina were infrequently detected (14.7%), likely reflecting the limited number of yeast-form species within this lineage (**Supplementary Fig. 2**).

The number of co-occurring yeast subphyla per sample varied considerably across environments (**Fig. 3a and Supplementary Fig. 3**). Anthropogenic and cropland habitats showed the highest yeast subphylum co-occurrences (**Fig. 3a**), often with three or more subphyla detected at 79.8% and 78.7%, respectively. Such human-influenced systems likely foster richer yeast assemblages by promoting multiple clades through management practices that enhance substrate heterogeneity and nutrient availability, increasing co-occurrence complexity (S. Zhu et al., 2023). Meta-analyses further show that organic amendments boost fungal diversity and shift community composition relative to mineral fertilization (Ma et al., 2018; Semenov, Krasnov, Semenov, & van Bruggen, 2022). In contrast, certain sample types (lichen, moss, root, and fungal sporocarp) frequently yielded only one or two yeast subphyla (only ∼45%–64% of these samples had more than one subphylum present), indicating more specialized or constrained communities (**Supplementary Fig. 3**). When restricting to samples where only a single yeast subphylum was detected, Agaricomycotina emerged as the most frequent lineage across most sample (**Supplementary Fig. 4**) and environment types (**Fig. 3b**). This pattern highlights their ecological versatility and suggests that members of this group are particularly well-suited to persist under conditions where other yeast lineages are absent or excluded. Together, these results reveal structured patterns of subphylum distribution and co-occurrence shaped by environmental complexity, niche specialization, and lineage-level ecological traits.

**Figure 3.**
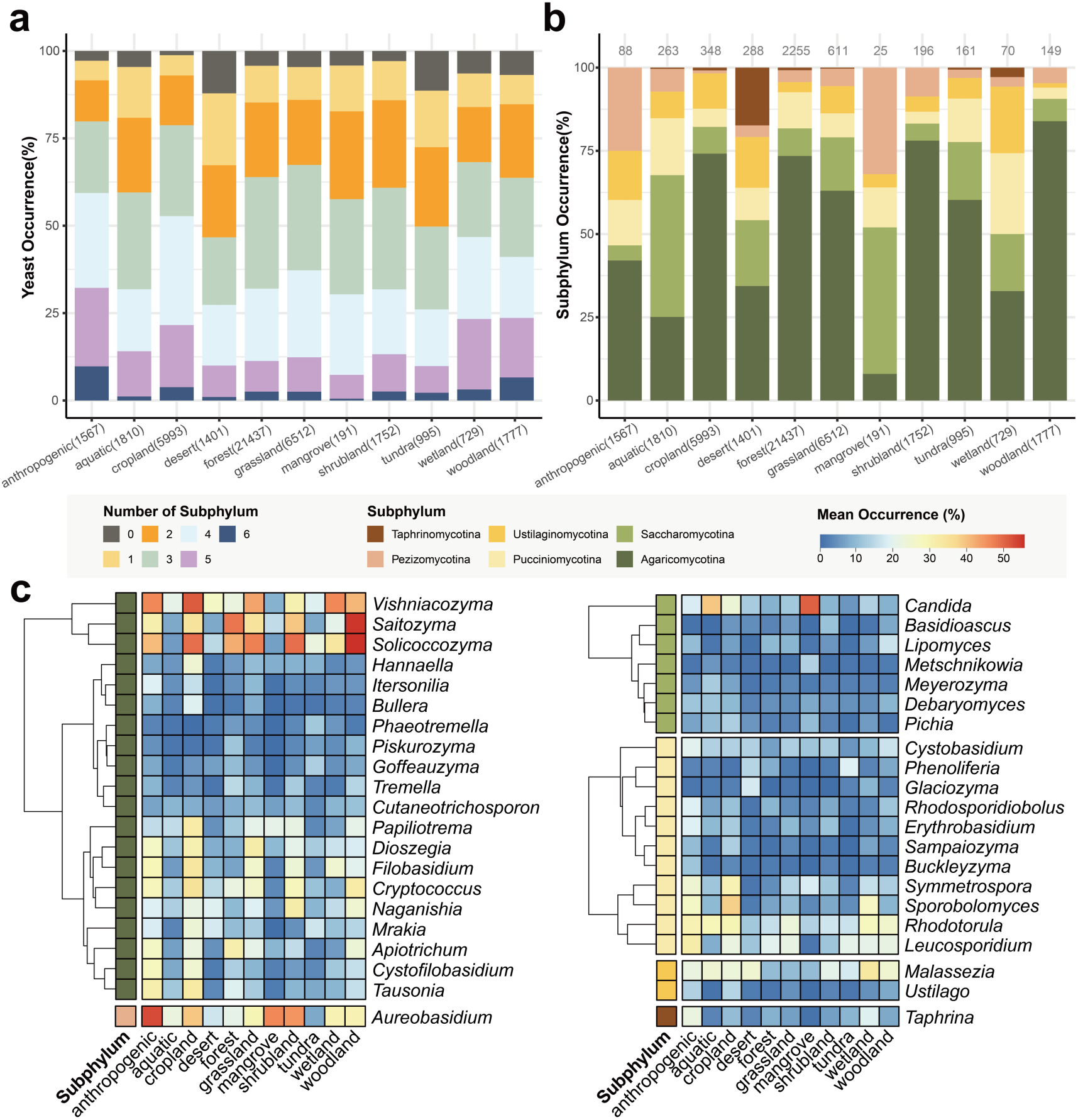
Habitat associations of yeast communities across environments. **(a)** Proportion of samples containing zero to six yeast subphyla across environment types. **(b)** Occurrence frequency of each yeast-forming subphylum across broad environments in samples where only a single subphylum was detected. The numbers above bars indicate the sample numbers included. **(c)** Heatmaps of genus-level yeast occurrence (>10% frequency) across environments, clustered by genus and colored by subphylum. Genera such as *Vishniacozyma*, *Solicoccozyma*, and *Saitozyma* were broadly distributed across environments, while others like *Candida*, *Pichia*, *Meyerozyma*, and *Debaryomyces* showed stronger habitat specificity, particularly within host-associated or anthropogenic niches.

To focus on ecologically meaningful patterns, our genus-level occurrence analysis was limited to taxa present in at least 10% of samples within one or more environments or sample types, thereby excluding rare or sporadic taxa and included a total of 42 genera (**Fig. 3c**). Within Agaricomycotina, *Vishniacozyma*, *Solicoccozyma*, and *Saitozyma* stood out as most broadly distributed and enriched in over 11.4% of samples across cropland, woodland and aquatic environments. Among Pucciniomycotina, *Rhodotorula*, *Sporobolomyces*, and *Leucosporidium* were prevalent across anthropogenic, cropland, and grassland settings, with occurrence rates generally exceeding 18.3%, suggesting their capacity to colonize and persist in disturbed or nutrient-enriched environments. Within Saccharomycotina, *Candida* stood out as particularly widespread, detected in over 17.7% of aquatic and anthropogenic habitats. Across the more filamentous yeast lineages (Ustilaginomycotina, Pezizomycotina, Taphrinomycotina), only four genera (*Malassezia, Taphrina, Ustilago, Aureobasidium*) exceeded the 10% occurrence threshold, and even these reached only moderate frequencies (∼7.7–52.3% of samples at most for *Aureobasidium*). These genus-specific occurrence patterns collectively highlight the ecological versatility of yeasts and reveal the diverse ecological strategies employed by yeast taxa in different global habitats.

To complement the environment-level heatmaps, we examined genera detected in at least one major sample category (frequency >10%; **Supplementary Fig. 5**). This analysis highlighted clear habitat preferences among yeast genera within and across subphyla. For example, *Malassezia* (Ustilaginomycotina) showed strong and specific occurrence in glacial ice debris and coral samples, consistent with recent findings of its presence in extreme or host-associated environments beyond human skin (Steinbach et al., 2023). Within Saccharomycotina, genera such as *Candida*, *Debaryomyces*, and *Cyberlindnera* displayed broad occurrence across various nutrient-rich or osmotolerant habitats, including aquatic environments, sediments, and decaying plant tissues. Other genera such as *Pichia* and *Meyerozyma* were largely confined to anthropogenic or nutrient-enriched environments. Pucciniomycotina genera such as *Rhodotorula*, *Sporobolomyces*, and *Leucosporidium* were observed across diverse environments, including shoot tissues, rhizosphere soils, and aquatic substrates. Agaricomycotina genera—including *Vishniacozyma*, *Solicoccozyma*, and *Tausonia*—were broadly distributed across topsoil, litter, and mosses, often co-occurring in forest and alpine habitats. Together, these genus-level occurrence patterns illustrate clear distinctions between generalist yeasts capable of colonizing diverse substrates, and specialist yeasts adapted to specific niches, underscoring consistent ecological preferences and reinforcing the structured nature of yeast community assembly.

### Low relative abundance of yeasts across environments

Having established the pervasive presence of yeasts across environments, we next examined their relative abundance within fungal communities (**Fig. 4a**). Among all subphyla, only Agaricomycotina reached notable relative abundances, with a median of 0.6% and values exceeding 10% in some samples (**Fig. 4b**). This numerical dominance likely reflects the diversities and ecological traits of Agaricomycotina, including broader environmental tolerances, efficient spore dispersal mechanisms, and competitive metabolic strategies that facilitate their persistence and proliferation across diverse habitats (Fell, Boekhout, Fonseca, Scorzetti, & Statzell-Tallman, 2000; Kurtzman & Boekhout, 2017). In contrast, other yeast subphyla such as Pucciniomycotina (0.02%), Saccharomycotina (0.01%), and especially Pezizomycotina, Taphrinomycotina, and Ustilaginomycotina (all ∼0%) occurred at substantially lower median abundances (**Fig. 4b**). When considering only the yeast component itself, Agaricomycotina consistently accounted for the largest share, with a median relative abundance of 84.9%, followed distantly by Pucciniomycotina (1.52%) and Saccharomycotina (0.38%) (**Fig. 4c**). At the genus level, a similar pattern of consistently low relative abundance was observed. Even the most frequently detected genera such as *Solicoccozyma*, *Saitozyma*, and *Vishniacozyma* generally accounted for <0.1% of total fungal reads within their respective environments, with the vast majority of genera remaining below 0.05% (**Supplementary Fig. 6**). These findings confirm the persistent rarity of yeasts in microbial communities, despite their global ubiquity, and underscore Agaricomycotina as the numerically dominant yeast lineage across diverse environments.

**Figure 4.**
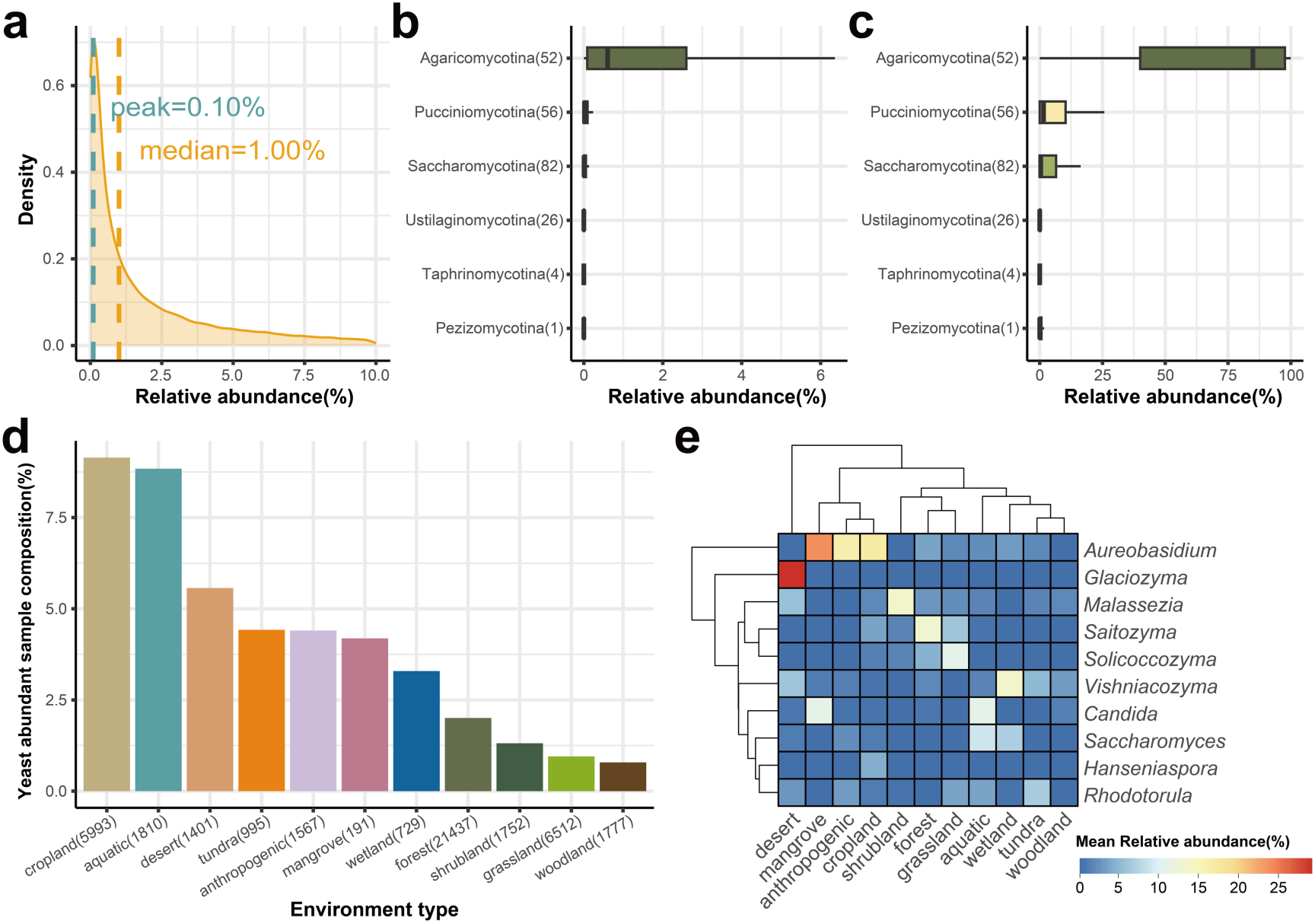
Relative abundance patterns and ecological contexts of yeast across samples. **(a)** Density distribution of yeast relative abundance (% of total fungal reads) across all 44,164 samples. **(b)** Relative abundance of yeast subphyla expressed as a percentage of all fungal reads. **(c)** Composition of the yeast community, showing the proportion of each subphylum within total yeast reads. **(d)** Distribution of yeast-rich samples (relative abundance > 25%) across broad environment types, expressed as a percentage of total samples within each environment. Numbers in brackets along the x-axis indicate the total number of samples of each environment type. **(e)** Heatmap of mean relative abundances (%) of the ten most abundant yeast genera across environment types.

### Exceptionally yeast-rich samples and their ecological contexts

Although yeasts typically constitute only a small fraction of microbial communities, we identified a notable subset of 1,460 samples out of 44,164 total (∼3.3%) in which yeasts exceeded 25% of total fungal reads. Environmentally, yeast-rich samples were not evenly distributed. Instead, they were most frequently found in cropland (9.1%), aquatic (8.8%), and desert (5.6%) environments, with progressively lower proportions in forest, grassland, and woodland habitats (**Fig. 4d**). These patterns suggest that highly disturbed regimes or extreme habitats may support episodic yeast proliferation. At the genus level, we further explored yeast-rich samples by identifying the ten genera with the highest mean relative abundance (**Fig. 4e**). Strikingly, eight of these genera—including *Aureobasidium, Glaciozyma, Malassezia, Saitozyma*, *Solicoccozyma*, *Vishniacozyma, Candida*, and *Rhodotorula*—were also among the most frequently occurring across the entire dataset, suggesting they are not only widespread but occasionally dominant. Despite this, their mean relative abundances within yeast-rich samples generally remained below 10%, with the exception of *Aureobasidium*, which reached >23% in a subset of samples. These values indicate that most yeast-rich samples are characterized by one or a few highly dominant OTUs, rather than a diverse assemblage of abundant yeast taxa. Interestingly, *Hanseniaspora* and *Saccharomyces*—taxa that are relatively rare across the global dataset—emerged among the top ten in relative abundance (**Fig. 4e**). These genera were particularly prominent in aquatic and cropland environments, such as fermenting substrates and nutrient-enriched waters (**Supplementary Fig. 7**, (Bickford, Zak, Kowalski, & Goldberg, 2020; Li et al., 2022; Talas, Stivrins, Veski, Tedersoo, & Kisand, 2021)). Their episodic dominance likely reflects their ecological strategy of rapid proliferation in sugar-rich, disturbed, or anaerobic conditions, consistent with their well-known roles in fermentation and gut-associated niches.

To further characterize these yeast-rich samples, we examined their beta diversity patterns relative to environmental and substrate type. Principal coordinate analysis (PCoA) based on Aitchison distances showed clear clustering associated primarily with habitat types, suggesting strong niche differentiation among these yeast-dominant communities (**Supplementary Fig 8**). Specifically, forest, woodland, shrubland, and grassland environments clustered distinctly from cropland and aquatic samples along the first principal component. Similarly, sample types showed evident separation, with soils, topsoils, and rhizosphere samples clustering separately from plant-associated samples (shoot, mosses) and wood substrates (deadwood). These diversity patterns reinforce the interpretation that yeast dominance in specific samples typically results from ecological filtering, where environmental and substrate-specific conditions select for a narrow range of highly opportunistic yeast taxa rather than a broad spectrum of diverse OTUs.

### Yeasts in metagenomes

To complement the amplicon analyses, we examined two large shotgun metagenomic datasets to test whether yeasts show comparable patterns when sequencing bias is removed. The first dataset comprised 750 samples from the subset of Earth Microbiome Project (EMP; (Shaffer et al., 2022)), spanning natural environments such as soils, aquatic systems, and animal hosts. The second contained 1,915 curated food-associated metagenomes (Carlino et al., 2024), representing a spectrum of fermented and unfermented substrates. We first benchmarked yeast detection in 125 EMP samples for which both ITS amplicon and metagenomic data were available. A k-mer–based classifier (Kraken2 + Bracken, (Lu, Breitwieser, Thielen, & Salzberg, 2017; Wood, Lu, & Langmead, 2019) consistently recovered more yeast-positive samples than the marker-gene approach EukDetect (Lind & Pollard, 2021), which is limited by database coverage (**Supplementary Fig 9**). Where species such as *Saccharomyces cerevisiae* were represented in its reference set, both tools performed comparably. Given its broader recall, Kraken2 + Bracken was used for all downstream analyses, with a confidence threshold of ≥ 0.4 and a minimum of 10,000 reads per sample to ensure robust assignments.

Across habitats, yeasts were detected in most categories but with marked contrasts in prevalence and detection thresholds (**Fig. 5**). In the EMP dataset, they were most common in host-associated niches—53.3% in the animal proximal gut, 39.2% in plant tissues, and 37.9% in the animal distal gut—while free-living environments showed substantially lower frequencies (**Fig 5a**). In the food metagenomes, yeast detection approached ubiquity, often reaching 100% in fermented products and >70% in probiotics (**Fig 5a**). Detection probability correlated strongly with sequencing depth (**Fig 5b**): in EMP samples, those with at least one detected yeast had a median of 2.0×10⁷ reads, roughly twice that of yeast-negative samples (1.1×10⁷). In contrast, yeast-rich food samples required fewer reads for confident detection, with medians around 5.4×10⁶ compared to 3.6×10⁷ in yeast-negative ones, consistent with their higher absolute yeast loads. Samples lacking any fungal reads were not necessarily low in sequencing depth—some exceeded 10⁸ total reads—highlighting that absence is often due to biological rarity or database incompleteness rather than insufficient coverage. Overall, samples with >10⁷ reads were generally sufficient to recover at least one yeast in EMP, whereas in the food dataset, yeasts were frequently detectable even below this threshold. When samples were classified as containing Yeast, Other Fungi, or None, the proportion lacking any fungal reads reached 91.6% in EMP but only 11.1% in food datasets, reflecting both genuine ecological scarcity and uneven sequencing depth across environments.

**Figure 5.**
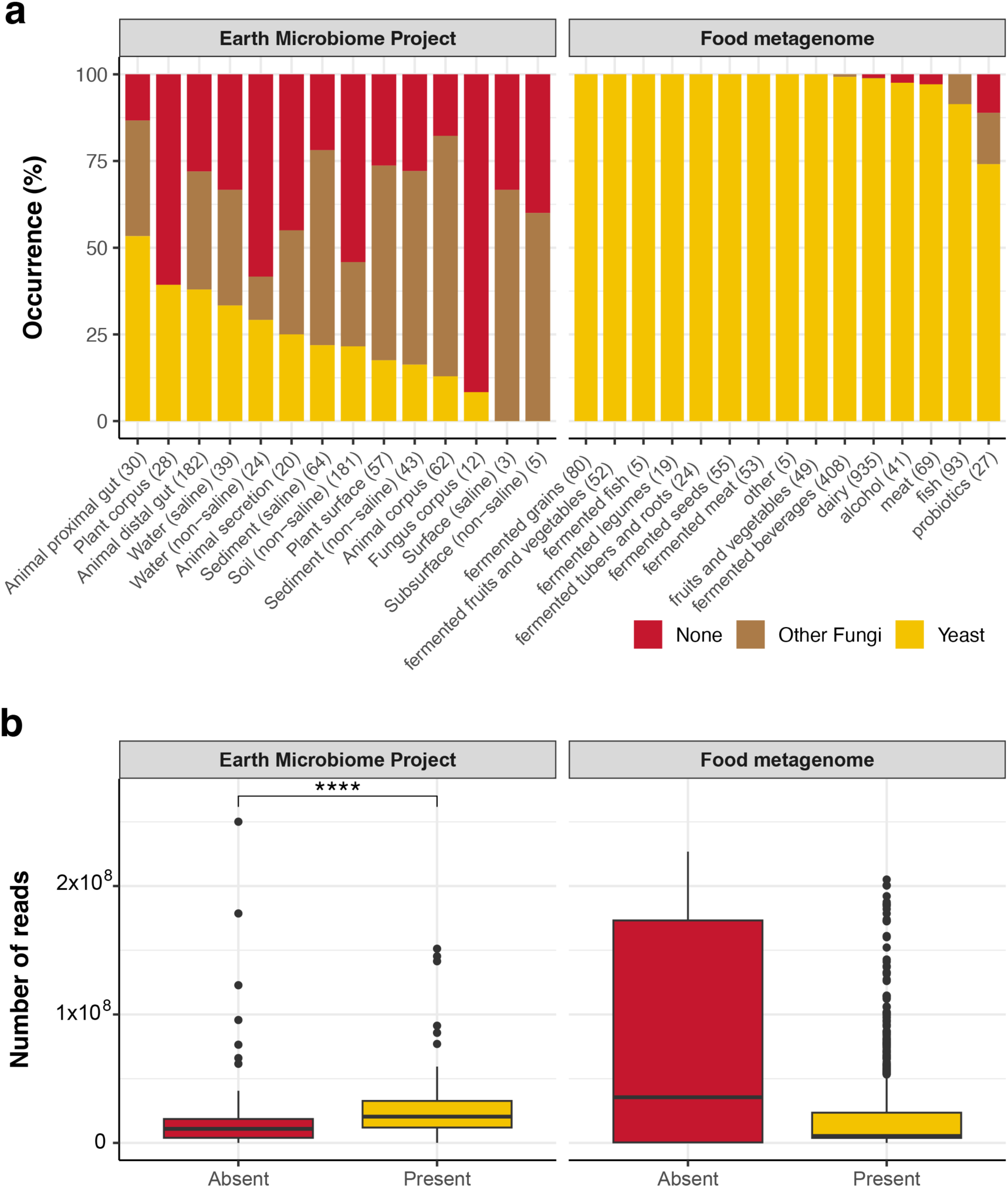
Prevalence and detection thresholds of yeasts across metagenomic datasets. **a.** Proportion of samples classified as containing yeasts (yellow), only non-yeast fungi (brown), or no fungal reads (red) in the EMP and food metagenomes. **b.** Relationship between sequencing depth and yeast detection for the same datasets.

Subphylum-level occurrence patterns revealed clear contrasts between wild and anthropogenic microbiomes, but Saccharomycotina emerged as the most frequently detected lineage in both datasets (**Fig. 6**). In the Earth Microbiome Project samples, Saccharomycotina occurred across nearly all categories but reached its highest frequencies in host-associated environments. In contrast, food metagenomes showed an even stronger dominance of Saccharomycotina detected on average 96.7% of samples amongst categories, particularly in fermented and probiotic samples, followed by moderate occurrences of Pezizomycotina and Pucciniomycotina. These patterns parallel the amplicon-based findings, highlighting that fermentative yeasts are ubiquitous across ecosystems but attain their greatest prevalence under nutrient-rich, anthropogenic conditions, whereas natural habitats harbour a more compositionally diverse assemblage of subphyla with lower and more variable frequencies.

**Figure 6.**
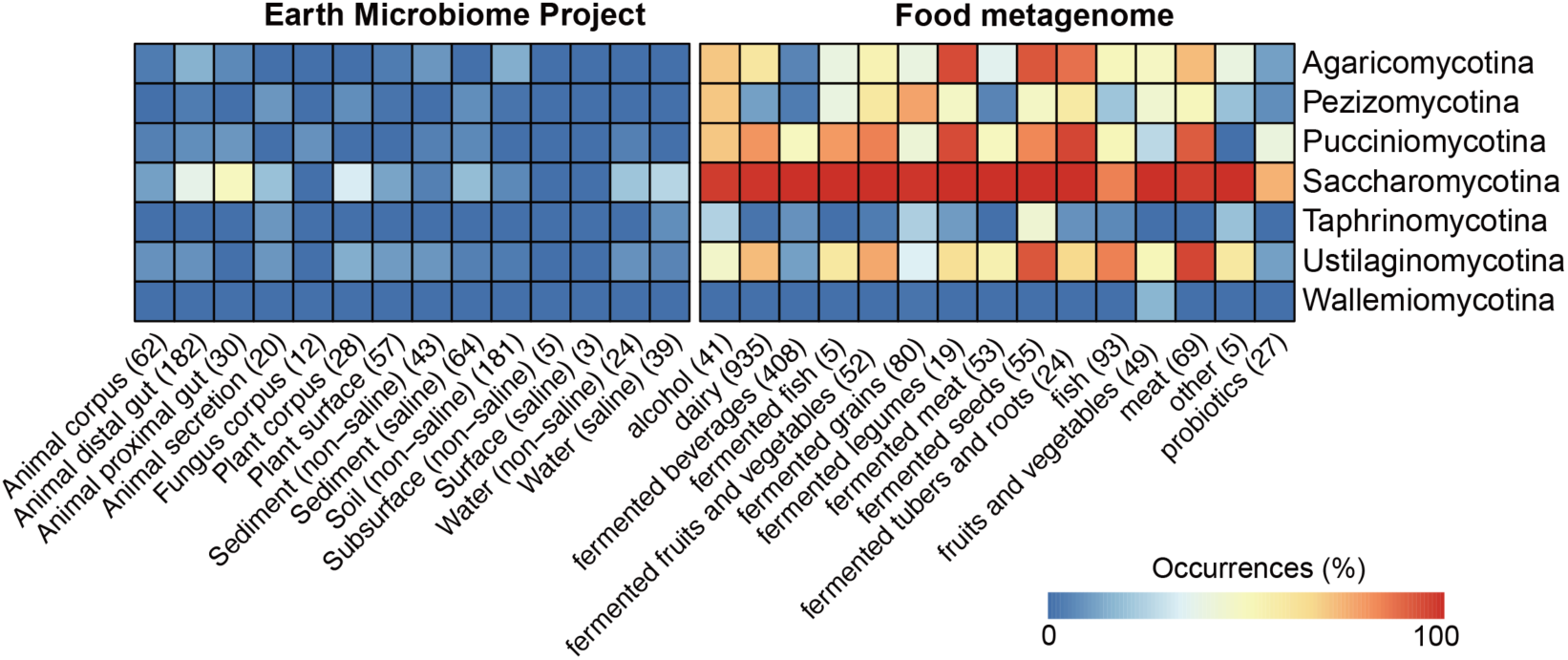
Occurrence frequency of fungal subphyla across metagenomic categories. Heatmap showing the percentage of samples in which each fungal subphylum was detected in the Earth Microbiome Project (left) and food metagenomes (right). Numbers in brackets denote number of samples in each category.

At the quantitative level, Saccharomycotina displayed the highest relative abundance in both metagenomic datasets, though its dominance was more apparent in human-related samples (**Fig. 7**). Median relative abundances, calculated as a proportion of all fungal reads, were 0.13% for Saccharomycotina and 0.12% for Pezizomycotina in EMP, compared with 1.32% and 0.57% in food metagenomes. Outliers in fermented beverages approached complete dominance by *Saccharomycotina* (∼100%), whereas the highest EMP values, around 70%, were contributed by Pezizomycotina. When analysis was restricted to yeast reads alone, *Saccharomycotina* remained the prevailing lineage in both datasets, indicating that most detectable yeast diversity is concentrated within this group. Despite these occurrences, fungal reads constituted only a minute fraction of total metagenomic sequences, with median unclassified proportions of 99.9% in EMP and 96.5% in food datasets. Yeasts themselves were typically rare (median 0% in EMP and 1.38% in food), underscoring the “curse of low abundance” that limits their recovery from shotgun data. Nonetheless, their consistent detection across diverse contexts—particularly in high-coverage or nutrient-rich samples—demonstrates that even rare fungal signals can be ecologically informative when viewed within multi-kingdom microbiomes.

**Figure 7.**
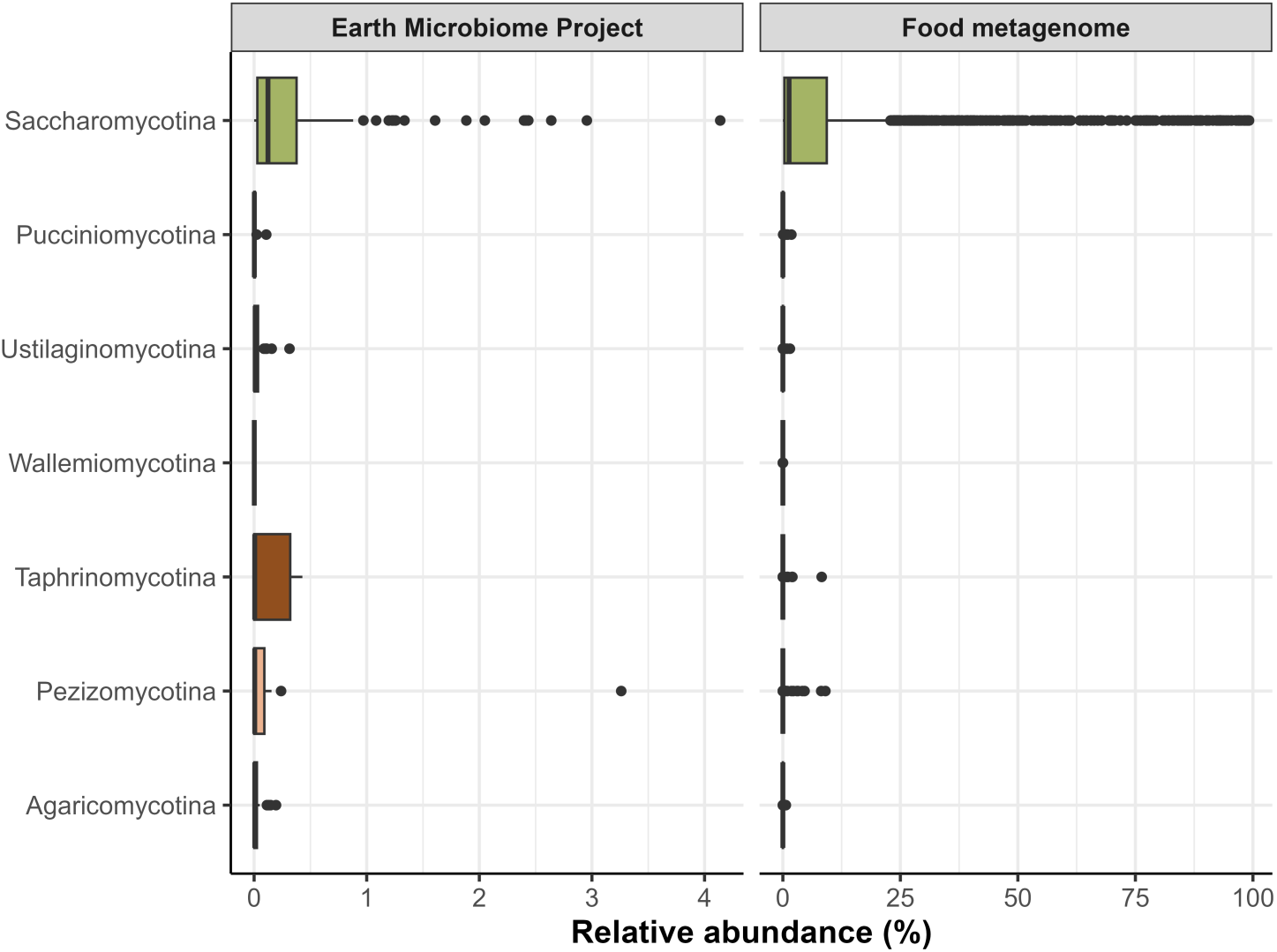
Relative abundance of fungal subphyla across metagenomic datasets. Boxplots show the distribution of relative abundances (%) of major fungal subphyla in the Earth Microbiome Project (left) and food metagenomes (right). Each point represents a sample, and abundances are calculated as proportions of total fungal reads.

## Discussion

Our global synthesis confirms that yeasts are widespread constituents of microbial communities across terrestrial, aquatic, and host-associated environments, yet they are consistently underrepresented in both detection and interpretation. Their distribution is not stochastic: yeast communities exhibit structured patterns shaped by lineage identity, environmental context, and substrate type. Basidiomycetous yeasts—especially within *Agaricomycotina* and *Pucciniomycotina*—were the most frequently detected across habitats, while *Saccharomycotina* were more narrowly associated with anthropogenic and aquatic settings. Although yeasts typically occur at low relative abundance, our analyses reveal that they can periodically dominate niches under specific ecological filters, including high-sugar substrates, saline environments, and disturbed or nutrient-enriched habitats. Our findings align with the ‘rare biosphere’ concept (Lynch & Neufeld, 2015), wherein organisms that are rare in abundance can still be widespread and ecologically important. The consistent habitat associations we observed indicate that yeast community assembly is governed more by niche filtering and habitat specificity than by random colonisation. This structured yet low-abundance presence underscores the need to re-evaluate assumptions of yeast rarity and ecological insignificance, recognising them instead as ecologically active and functionally important players in microbial assemblages.

While yeasts are usually rare, certain ecological contexts consistently favour their dominance. Beyond the classic examples of *Saccharomyces* in fermented foods and *Malassezia* on human skin, other genera became locally abundant under specific filters. In sugar-rich substrates, *Metschnikowia* dominates flower nectar, ripe fruits, and fermentation niches due to its rapid fermentative metabolism and tolerance of acidity (Dhami, Hartwig, & Fukami, 2016; Pozo, Lachance, & Herrera, 2012). In saline environments, *Debaryomyces hansenii* thrives in coastal waters, salt marshes, and hypersaline habitats owing to its exceptional osmotolerance (Navarrete, Estrada, & Martínez, 2022). Cold-adapted yeasts such as *Naganishia* and *Glaciozyma* are enriched in polar soils, glacial ice, and alpine lakes, where their psychrophilic physiology gives them a competitive edge (Buzzini, Turk, Perini, Turchetti, & Gunde-Cimerman, 2017; Segal-Kischinevzky et al., 2022). Our analysis also identified consistent enrichment of *Pichia* and *Meyerozyma* in disturbed, nutrient-enriched orchard soils (S. Zhu et al., 2023). Together, these findings show that yeast dominance, though episodic, is not confined to fermentation or host-associated niches but arises broadly under conditions of disturbance, enrichment, or environmental stress.

A central implication of our findings is that rarity in abundance does not equate to ecological insignificance. Even at low relative abundance, yeasts contribute disproportionately to ecosystem processes. In soils, yeasts solubilise phosphate, degrade organic matter, sequester heavy metals, and act as biocontrol agents against pathogens (S. Zhu et al., 2023). Some soil yeasts even promote plant growth by enhancing mycorrhizal colonization or producing beneficial secondary metabolites (Yurkov, 2018). In aquatic systems, reviews emphasise that planktonic fungi, including yeasts, are integral to nutrient cycling and carbon flux, with fungal biomass at times approaching bacterial levels in coastal waters (Sen, Sen, & Wang, 2022). Host-associated environments show similar dynamics: despite comprising only 0.01–0.1% of gut microbes, fungi can shape host immunity and microbiota composition. Liu et al. (2025) highlight how intestinal fungi engage in antagonistic and synergistic interactions with bacteria, influencing colonisation resistance, barrier function, and immune modulation even at low abundance (Liu et al., 2025). Large-scale analyses further reveal that human gut mycobiomes form stable enterotypes (dominated by *Candida*, *Saccharomyces,* or *Aspergillus*) that correlate with host age, metabolic state, and disease risk, underscoring structured contributions despite numerical rarity (Lai et al., 2023). These observations echo the broader recognition that microbial impact is not proportional to read counts. Systematic exclusion of yeasts from microbiome analyses therefore risks overlooking functionally important guilds.

This perspective is increasingly supported by recent reviews and empirical studies. For example, Rosa et al. (C. A. Rosa et al., 2023) emphasized the diversity and ecological relevance of tropical forest yeasts, while Sen et al. (Sen et al., 2022) reviewed the pivotal but underexplored role of marine yeasts in planktonic ecosystems. These perspectives reinforce our findings and illustrate a growing recognition that yeasts—often treated as peripheral—should be embedded within core microbiome frameworks. One genus that exemplifies an ‘overlooked yet important’ yeast is *Aureobasidium*, which we found at both high occurrence and relative abundance across diverse environments. This cosmopolitan “black yeast” has long been known to colonize rock surfaces, leaf litter, and bark (Urzì, De Leo, Lo Passo, & Criseo, 1999), and has even been proposed as a model for ecological versatility due to its polyextremotolerance with biotechnological potential (Chi et al., 2009; Xiao et al., 2024). Our meta-analysis reaffirms these patterns and highlights *Aureobasidium*’s status as a persistent and sometimes dominant taxon. This theme extends beyond *Aureobasidium*: a recent metagenomic survey of skin microbiomes detected an abundant presence of black yeast-like fungi (Chaetothyriales), including *Exophiala* and *Cyphellophora*, even in healthy individuals (Voidaleski, Costa, de Hoog, Gomes, & Vicente, 2023).

Future research on yeasts in microbiomes should prioritize inclusive, multi-kingdom approaches that enable robust detection and ecological interpretation. Amplicon sequencing using ITS regions remains essential for fungal profiling, but improved primer design is needed to reduce taxonomic bias against lineages such as *Saccharomycetales* and *Malasseziales* (Leho Tedersoo et al., 2015). Alternative methods such as shotgun metagenomics or long-read sequencing (e.g. PacBio or Oxford Nanopore) offer the advantage of bypassing primer bias while providing higher taxonomic resolution and insights into functional potential (D’Andreano, Cusco, & Francino, 2021; Renzi et al., 2024; L. Tedersoo et al., 2022), but their utility is still constrained by poorly curated fungal genome databases and non-standardised workflows, leaving many reads unclassified (Renzi et al., 2024). In addition, protocols for DNA extraction should be optimized for fungal cell walls, as under-recovery of yeast DNA can skew abundance estimates (Aziz et al., 2015). The integration of quantitative tools—qPCR, flow cytometry, and stable isotope probing—can help estimate yeast biomass and activity. Cultivation remains indispensable for discovering novel metabolism in underexplored taxa such as *Tausonia* and *Metschnikowia*, and co-culture experiments with bacteria, filamentous fungi, and plant hosts are vital to reveal ecological outcomes ranging from facilitation to antagonism (Frey-Klett et al., 2011). Together, these methodological and conceptual advances will bring yeast ecology onto equal footing with bacterial microbiome science.

In conclusion, this study reveals that yeasts, though often numerically minor constituents, are structured, widespread, and ecologically significant members of microbial communities across diverse ecosystems. By merging large-scale dataset analyses with detailed taxonomic profiling and literature reviews, we show that yeast diversity reflects distinct patterns shaped by environmental conditions, substrate availability, and lineage-specific traits. Notably, recent global surveys have uncovered yeasts in unexpected environments—such as the detection of *Malassezia* in marine ecosystems (Steinbach et al., 2023), highlighting how multi-kingdom microbiome studies continue to reveal overlooked fungal lineages. However, interpreting such findings requires caution: a recent analysis of marine metagenomes showed that *Malassezia* reads were strongly correlated with human-associated DNA, suggesting that many detections may result from contamination rather than true environmental colonization (Rahimlou, Amend, James, & Taylor, 2025). Nevertheless, whether transient or resident, the recurring presence of yeasts across ecosystems underscores their ecological relevance. Embracing yeast diversity within comprehensive microbiome frameworks will refine community-level interpretations and help uncover the broader functional contributions of fungi in global ecosystems.

## Methods

### Selection of representative metabarcoding studies

To capture how yeasts have been represented in previous microbiome research, we assembled a dataset of 39 representative studies spanning terrestrial, aquatic, host-associated, and anthropogenic environments. Candidate publications were identified through keyword searches in Google Scholar using combinations of “yeast,” “microbiome,” “mycobiome,” “ITS,” “amplicons,” “metabarcoding,” and “metagenomics.” From the resulting pool, we prioritised studies that (i) explicitly reported fungal or yeast taxa in community surveys, (ii) employed high-throughput sequencing of fungal barcodes (ITS1, ITS2, or LSU) or shotgun metagenomics, and (iii) spanned multiple host or environmental contexts. Each study was reviewed to determine which yeast genera were mentioned, detected, or reported as dominant. Metadata extracted included sequencing method, target marker, and sample type. This curation is detailed in **Supplementary Table 1** and visualized in **Figure 1** and **Supplementary Fig 1**.

### Analysis of GlobalFungi dataset

We obtained the genus-level amplicon data and associated metadata from the GlobalFungi Database, with ITS1 and ITS2 regions included (Větrovský et al., 2020). Taxonomic information for each genus was assigned using R package *taxonomizr* (v0.11.1; https://github.com/sherrillmix/taxonomizr) and updated using package *rgbif* (v3.8.0; (Chamberlain & Boettiger, 2017). To identify yeast taxa, all genera were queried against The Yeasts Trust Database (https://theyeasts.org/) using a custom *selenium* crawler (v4.29.0; https://github.com/SeleniumHQ/selenium). Only genera confirmed to be listed in the database were classified as yeasts in downstream analyses. All analyses were conducted in the R environment (v4.3.3; https://www.R-project.org/). Amplicon datasets with fewer than 10,000 reads were removed using *phyloseq* (v1.46.0; (McMurdie & Holmes, 2013)). Data visualization was carried out with *ggplot2* (v3.5.2; (Villanueva & Chen, 2019)). The yeast occurrence map was generated using *ggspatial* (v1.1.9; https://CRAN.R-project.org/package=ggspatial), sf (v1.0.21; https://github.com/r-spatial/sf), *metR* (v0.18.1; https://github.com/eliocamp/metR), and *ggrepel* (v0.9.6; https://github.com/eliocamp/metR). Heatmaps were created using *pheatmap* (v1.0.13; https://github.com/raivokolde/pheatmap).

### Metagenomics

We analysed two public shotgun metagenomic datasets to complement the amplicon analyses: the Earth Microbiome Project (EMP; (Shaffer et al., 2022; Thompson et al., 2017)) and the curated food metagenomes (Carlino et al., 2024). Raw sequence data and sample metadata were downloaded from the European Nucleotide Archive (ENA) and Qiita (Gonzalez et al., 2018), respectively. After read cleaning, deduplication, and removal of host-derived sequences, we retained 750 EMP and 1,915 food-associated samples for downstream analyses. For EMP, 582 ITS metabarcoding samples and 125 paired ITS–metagenome samples were used for benchmarking detection performance. Taxonomic classification was performed using *EukDetect* (v1.3, (Lind & Pollard, 2021)) and *Kraken2 + Bracken* (v2.1.6 (Wood et al., 2019)/ v3.0.1 (Wood et al., 2019)), representing marker-gene and k-mer–based approaches, respectively. A fungal-only database comprising 14,616 curated genomes (hybrids removed and *Saccharomyces chiloensis* added) was used for the *Kraken2 + Bracken* pipeline, which demonstrated higher recall and was adopted for all subsequent analyses. Classifications were filtered with a confidence threshold ≥ 0.4 and restricted to samples containing ≥ 10,000 reads to ensure reliability. For ITS data, we applied stricter inclusion criteria (≥ 1,000 reads per sample; genus detections requiring ≥ 5% of reads or ≥ 20 reads) to minimise false positives. Relative abundances were computed at subphylum and genus levels using *phyloseq* (v1.46.0) in R, and visualisations were produced with *ggplot2* and *pheatmap*.

## Acknowledgement

Sample processing, sequencing, and core amplicon data analysis were performed by the Earth Microbiome Project (www.earthmicrobiome.org), and all amplicon sequence data and metadata have been made public through the EMP data portal.

## Author contributions

IJT and GL conceived the study. IJT and CPL performed the literature review. CPL analysed the data from Globalfungi. AG and NS analysed the metagenomics dataset. IJT and GL wrote the manuscript with input from all authors. All authors reviewed and approved the final version of the manuscript.

