## Supplementary Information for "A global synthesis of yeast in microbiomes"

**Supplementary Fig. 1. Full heatmap of yeast genera across published studies.** All yeast genera reported across 40 microbiome studies, with colours indicating whether each genus was mentioned (green), prevalent (yellow), or dominant (red)**.**

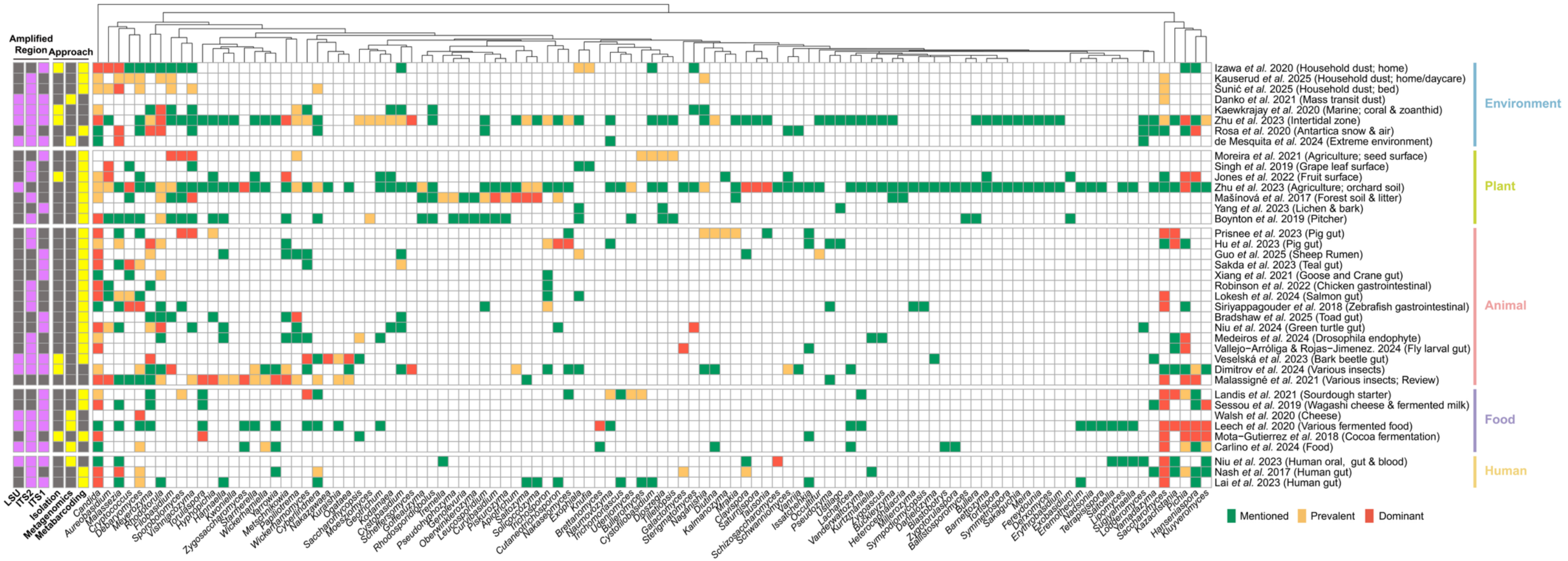

**Supplementary Fig. 2. Yeast-forming subphyla occurrence across environment types.** Colors represent whether the yeast is present (yellow) or absent (red). The numbers in brackets indicate the number of the environment type.

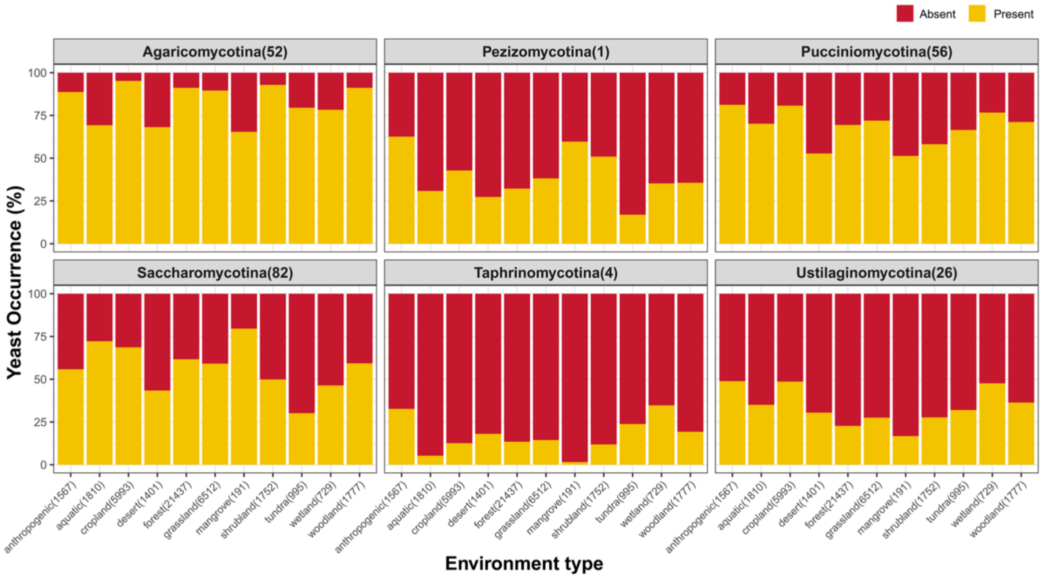


**Supplementary Fig. 3. Proportion of yeast subphyla co-occurrence across sample types.** The colour indicates the number of co-occurrence subphyla (zero to six). Numbers in the brackets in the x-axis represent the number of samples of each sample type.

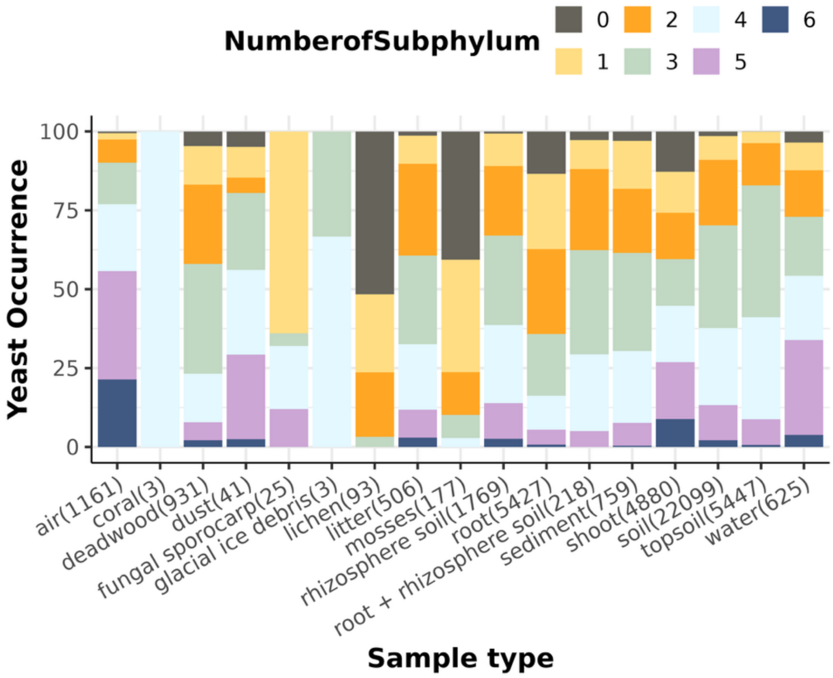

**Supplementary Fig. 4. Occurrence frequency of yeast-forming subphyla in single-subphylum samples across sample types.** The numbers above bars indicate the number of single-subphylum samples.

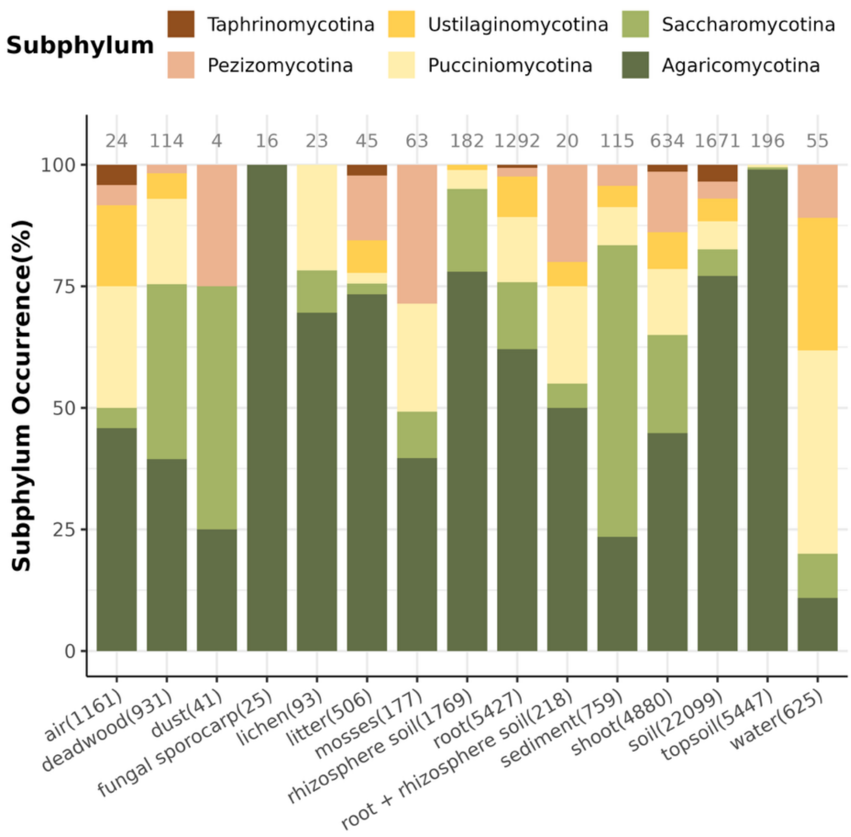

**Supplementary Fig. 5. Heatmap of genus-level yeast occurrence across sample type.** The cells are clustered by subphylum. Genera exhibiting less than 10% occurrence in all sample types were excluded. Cell colors represent the average percentage of occurrence for each genus, with genera clustered and annotated by subphylum.

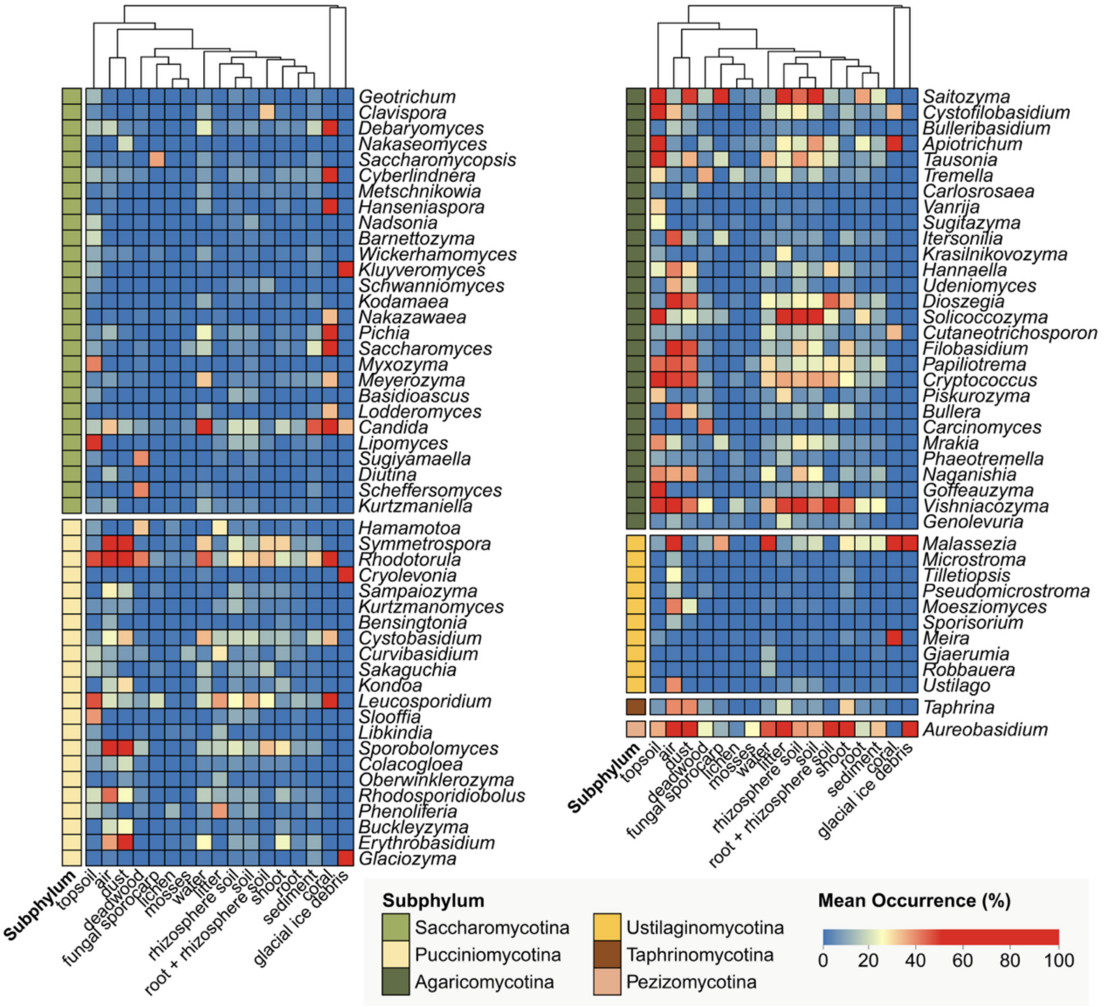

**Supplementary Fig. 6. Heatmap of relative abundance of top three abundant yeast genus of each subphylum across environment type.** The cell colours denote the average of relative abundance (%) within the environment type.

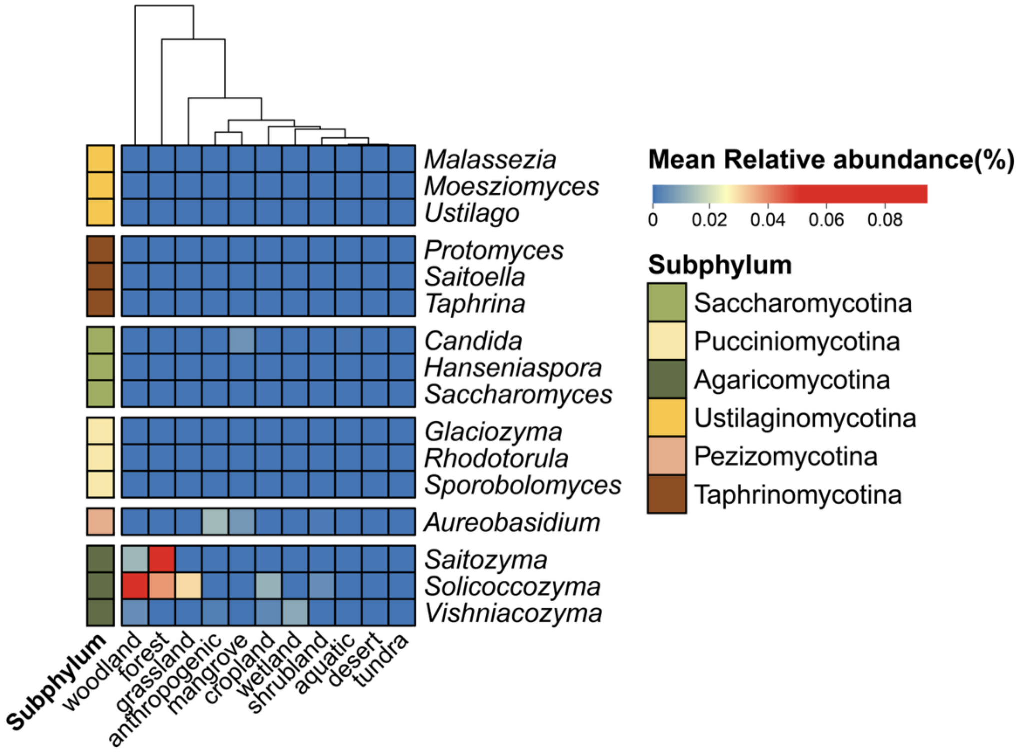

**Supplementary Fig. 7. Predominance of yeast genus *Hanseniaspora* and *Saccharomyces* in aquatic and cropland environments.** The dot color represents the ecosystem of the sample.

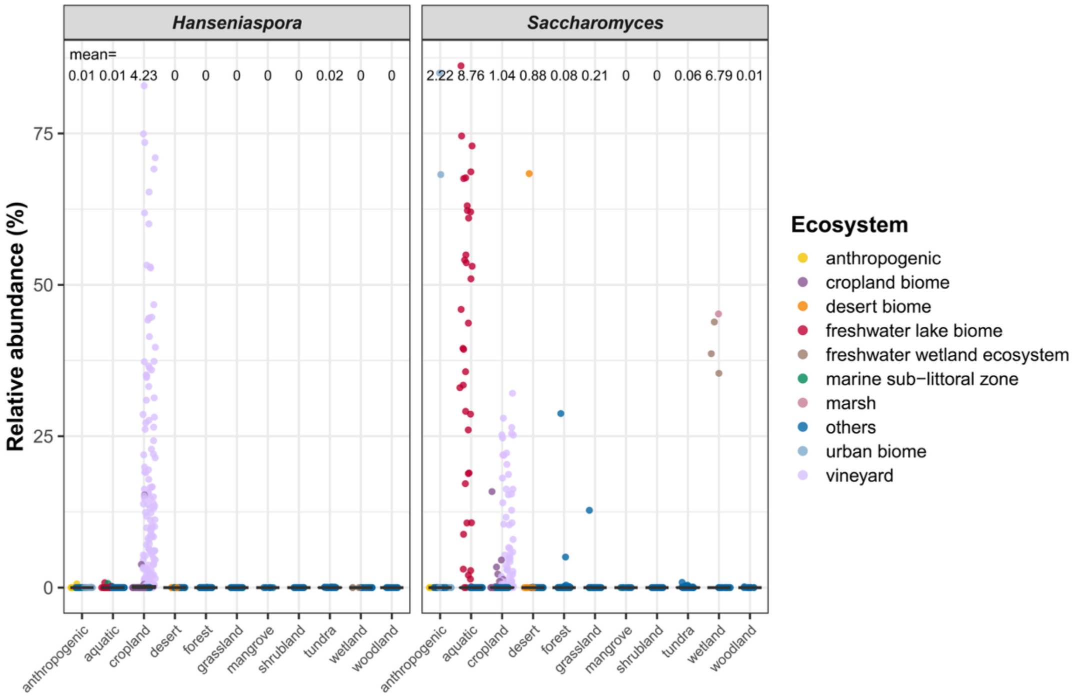

**Supplementary Fig. 8. Fungal composition of yeast-enriched samples across environment and sample types.** Samples with yeast relative abundance exceeding 25% were defined as yeast-enriched samples (1,460 out of 44,164 selected). The plots were calculated using the Aitchison distance and ordinated via principal coordinates analysis (PCoA), capturing 26.9% of the total variance. The dot colors represent environment type (left panel) and sample type (right panel), respectively.

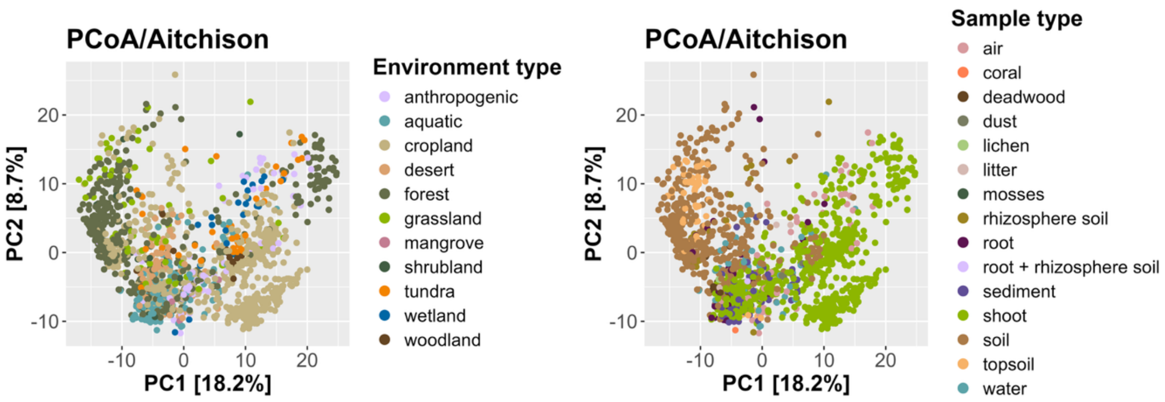

**Supplementary Figure 9. Comparison of yeast detection across metagenomic tools and environments.** (a–d) Earth Microbiome Project (EMP) samples; (e–h) food metagenomes. Panels (a, e) show the percentage of samples with detected yeasts using three approaches—ITS amplicon (grey), marker-gene (*EukDetect*; green), and k-mer (Kraken2 + Bracken; blue). Across nearly all categories, Kraken2 + Bracken recovered more yeast-positive samples than other methods. Panels (b, f) display the proportion of samples containing yeasts (yellow), only non-yeast fungi (brown), or no fungal reads (red). Panels (c, g) show the absolute abundances of the ten most represented yeast genera per category, while (d, h) highlight the dominance of *Saccharomyces* in selected animal and food-associated samples. Together, these analyses demonstrate that k-mer-based classification provides higher recall and greater sensitivity for yeast detection across both natural and anthropogenic microbiomes.

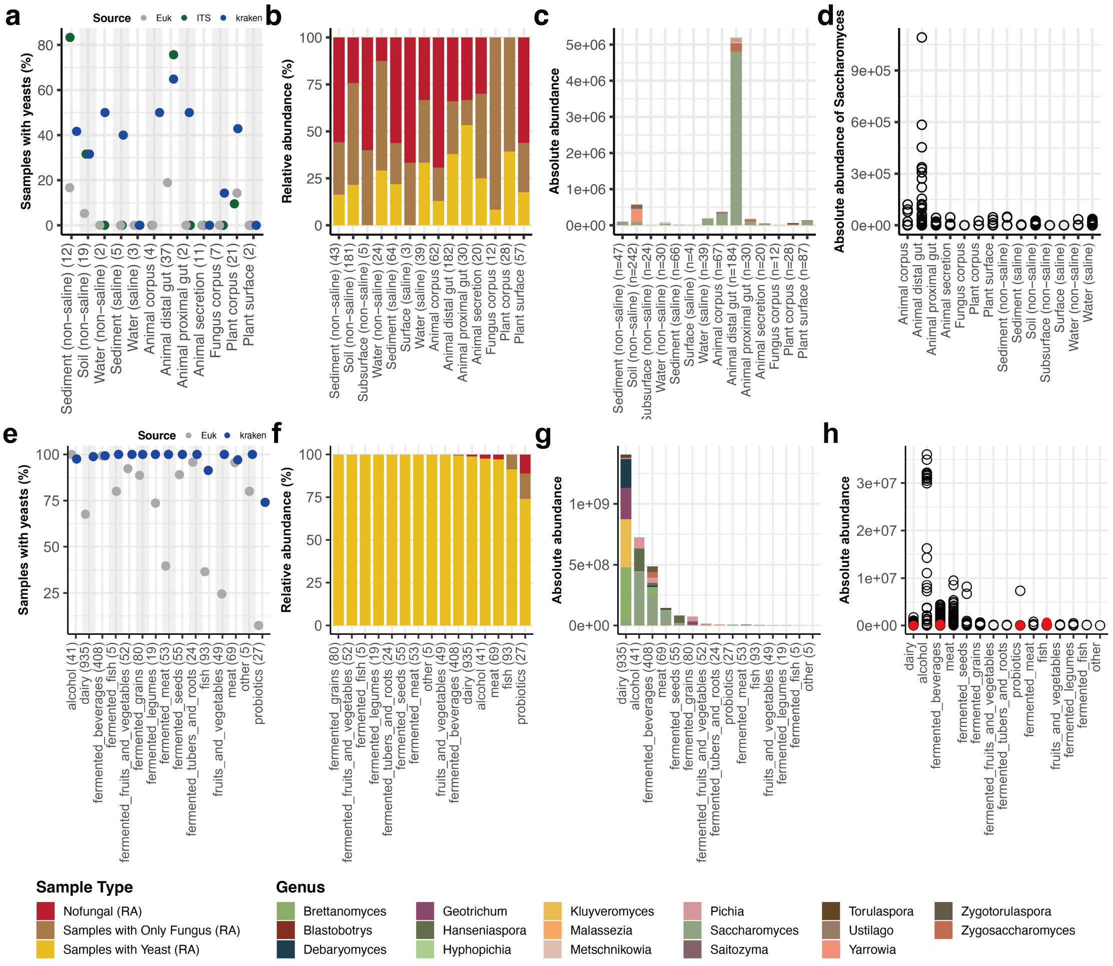
